# CellColoc: A modular, open-source workflow for cell colocalization, segmentation, and feature extraction in microscopy images

**DOI:** 10.64898/2026.07.26.740771

**Authors:** Fabrizio Musacchio, Henrike Antony, Arush Baijal, Denise Marie Hoffmann, Felix Christopher Nebeling, Sophie Crux, Martin Fuhrmann

## Abstract

Quantitative cell colocalization in fluorescence microscopy often depends on ad hoc combinations of image loading, segmentation, region selection, manual inspection, and spreadsheet post-processing. Such workflows are difficult to transfer across projects and often obscure how intermediate results were produced. We present CellColoc, an open-source Python workflow pipeline for segmentation-based cell colocalization, single-channel segmentation, and cell feature extraction in 2D and 3D microscopy images. CellColoc provides a modular workflow layer that integrates existing segmentation backends, including Cellpose and threshold-based methods, into reusable, script-driven analyses. The package supports channel-wise backend selection, interactive or reusable regions of interest, optional third-channel occupancy and cell-positivity analysis, z-cropping and z-projection, cached post hoc refinement of Cellpose thresholds, and reanalysis after manual mask editing. Analyses are executed from concise user scripts while reusable functionality is kept in a core package. Intermediate artifacts such as ROI masks, per-channel label masks, positive-cell masks, and structured result tables are written to a standardized results directory, promoting transparent inspection, reproducibility, and FAIR-aligned reuse. Public example datasets, a synthetic benchmark, and archived software releases accompany the package. By separating reusable analysis logic from project-specific configuration, CellColoc offers an extensible foundation for community-driven microscopy workflows that need transparent per-cell overlap classification, morphology readouts, and reusable batch analysis.

## Introduction

Open-source bioimage analysis has transformed microscopy research by making image processing, visualization, and quantification broadly reusable across laboratories [1, 2, 3]. Yet many recurring analyses in small and medium-sized projects are still operationalized as local combinations of viewers, copied scripts, hand-tuned segmentation settings, and spreadsheet post-processing. This is especially common for cell-focused fluorescence experiments in which users need to determine whether segmented cells are positive for markers in other channels, inspect region-specific subsets, and preserve intermediate masks for later quality control.

Colocalization analysis is particularly sensitive to workflow design because the biological question is often object-centric rather than pixel-centric. Guidance on quantitative colocalization has long emphasized that overlap measurements must be matched to the spatial scale of the hypothesis, the quality of segmentation, and the treatment of background and thresholds [4]. In many everyday applications, the primary outcome is a per-cell classification problem: does a segmented cell overlap sufficiently with a marker-defined structure to be called positive?

CellColoc addresses this practical gap as a reusable, open-source workflow pipeline for segmentation-based cell colocalization, single-channel segmentation, and feature extraction in 2D and 3D microscopy data. The package is designed around small user-editable scripts, explicit configuration objects, saved intermediate artifacts, and standardized result folders. In this way, it supports transparent, reproducible, and FAIR-aligned analysis practices [5, 6, 7, 8] while remaining modular enough to invite community extension.

## Related works

CellColoc sits within, and intentionally depends on, a much broader bioimage-analysis ecosystem. It is best understood as a workflow layer that bridges segmentation backends, interactive inspection, and reproducible export. Table 1 compares representative tools by their primary emphasis rather than by the union of every possible plugin or custom extension. It is important to make this distinction because several mature platforms can be extended far beyond their default use cases, and CellColoc is meant to complement rather than replace them. Commercial microscopy-analysis environments also exist, but we focus the comparison on tools with citable public technical descriptions and public software releases. We do this with intention in order to address the question, how CellColoc relates to inspectable, scriptable, community-facing analysis ecosystems rather than to every available end-user product.

**Table 1.** Representative position of CellColoc within the microscopy-analysis ecosystem. All listed tools are open-source and have public software releases. The comparison reflects the primary documented emphasis of each tool rather than every possible plugin or bespoke extension.

| Tool | Primary emphasis | Scripting | 3D | Per-cell object-based positivity |
| --- | --- | --- | --- | --- |
| Fiji / ImageJ [2, 3] | General bioimage analysis with plugins and macros | Yes | Yes | Partial |
| CellProfiler [9, 10, 11] | Modular pipelines and quantitative measurements | Partial | Yes | Partial |
| ilastik [12] | Interactive machine-learning-based segmentation and classification | Partial | Yes | No |
| QuPath [13] | Object-based 2D pathology and fluorescence analysis | Yes | Limited | Partial |
| Cellpose [14] | Segmentation backend | Yes | Yes | No |
| CellColoc | Script-driven microscopy workflow pipeline | Yes | Yes | Yes |

“Partial” indicates that the capability can typically be assembled through scripting, plugins, or composed modules but is not the main end-to-end workflow emphasis in the cited source. “Limited” indicates restricted or non-primary support relative to the other rows. To keep the comparison inspectable, we restrict it to tools with citable public descriptions and do not attempt to exhaust all commercial or plugin-specific variants.

The table distinguishes several neighboring tool classes that CellColoc is meant to complement. Fiji/ImageJ and QuPath are broad interactive analysis environments in which colocalization can be assembled as part of larger workflows [2, 3, 13]. CellProfiler emphasizes modular measurement pipelines [9, 10, 11], whereas ilastik and Cellpose primarily contribute segmentation or classification components rather than a complete end-to-end overlap-analysis workflow [12, 14]. CellColoc is positioned differently: it links microscopy-native I/O, per-channel segmentation choices, region of interest (ROI) handling, positivity calls, and persistent export into one compact scripted analysis pattern. In that sense, it functions less as a monolithic platform and more as a workflow layer that can sit alongside, or downstream of, several of the tools listed in Table 1.

The comparison suggests that CellColoc occupies a workflow-coordination layer within the ecosystem. Its main contribution is to package common cell-centered microscopy tasks into an explicit, small-script workflow in which the choice of segmentation backend, ROI strategy, and export structure remains visible and modifiable. This makes CellColoc particularly compatible with community contributions that add new dataset-specific scripts, additional segmentation backends, or richer downstream summaries without changing the overall analysis pattern.

## Methods

CellColoc addresses a workflow-engineering problem that recurs across many microscopy projects: file formats differ, segmentation backends require dataset-specific tuning, ROIs are often defined ad hoc, and quantitative calls must remain traceable after later refinement. The package is therefore built around a small set of explicit design principles that connect the software architecture to the analysis semantics and export model summarized in Box 1.

### Box1: Design goals of *CellColoc*

*CellColoc* was built around five explicit design goals:

**1. A small, inspectable workflow surface**. Routine analyses should remain centered on short Python scripts and a limited number of stable package entry points, so that project-specific channel roles, paths, and thresholds stay legible instead of being hidden in opaque project files.

**2. Explicit per-channel semantics and decision rules**. Every analysis branch should make channel assignment, segmentation backend choice, ROI handling, and positivity thresholds explicit, so that biological interpretation remains tied to visible analysis assumptions.

**3. Reproducibility by persistent intermediates**. Intermediate artifacts such as ROI masks, label masks, positive-cell masks, overlap tables, and refinement caches should be written to a predictable results structure so that runs can be audited, refined, or re-analyzed.

**4. Enabling microscopy-stack interoperability**. File-format heterogeneity should be delegated to dedicated microscopy I/O handlers so that downstream analysis can operate on consistent stack representations while preserving metadata access and explicit provenance.

**5. Modularity, extensibility, and community contribution**. New dataset-specific scripts, segmentation backends, postfilters, or summary exports should be easy to add without breaking the overall workflow model.

### Architecture and control flow

CellColoc addresses design Goals 1, 3, and 5 from Box 1 by separating reusable package logic from project-specific user scripts. The core package exposes modules for configuration, image I/O, ROI handling, segmentation, analysis, visualization, runtime helpers, and a parallel single-channel branch. End users typically do not edit internal package logic for routine analyses. Instead they create or adapt concise scripts that define dataset paths, channel roles, segmentation parameters, ROI behavior, and runtime settings before calling exported pipeline functions. The resulting arrangement is summarized in Figure 1 a,b.

**Figure 1:**
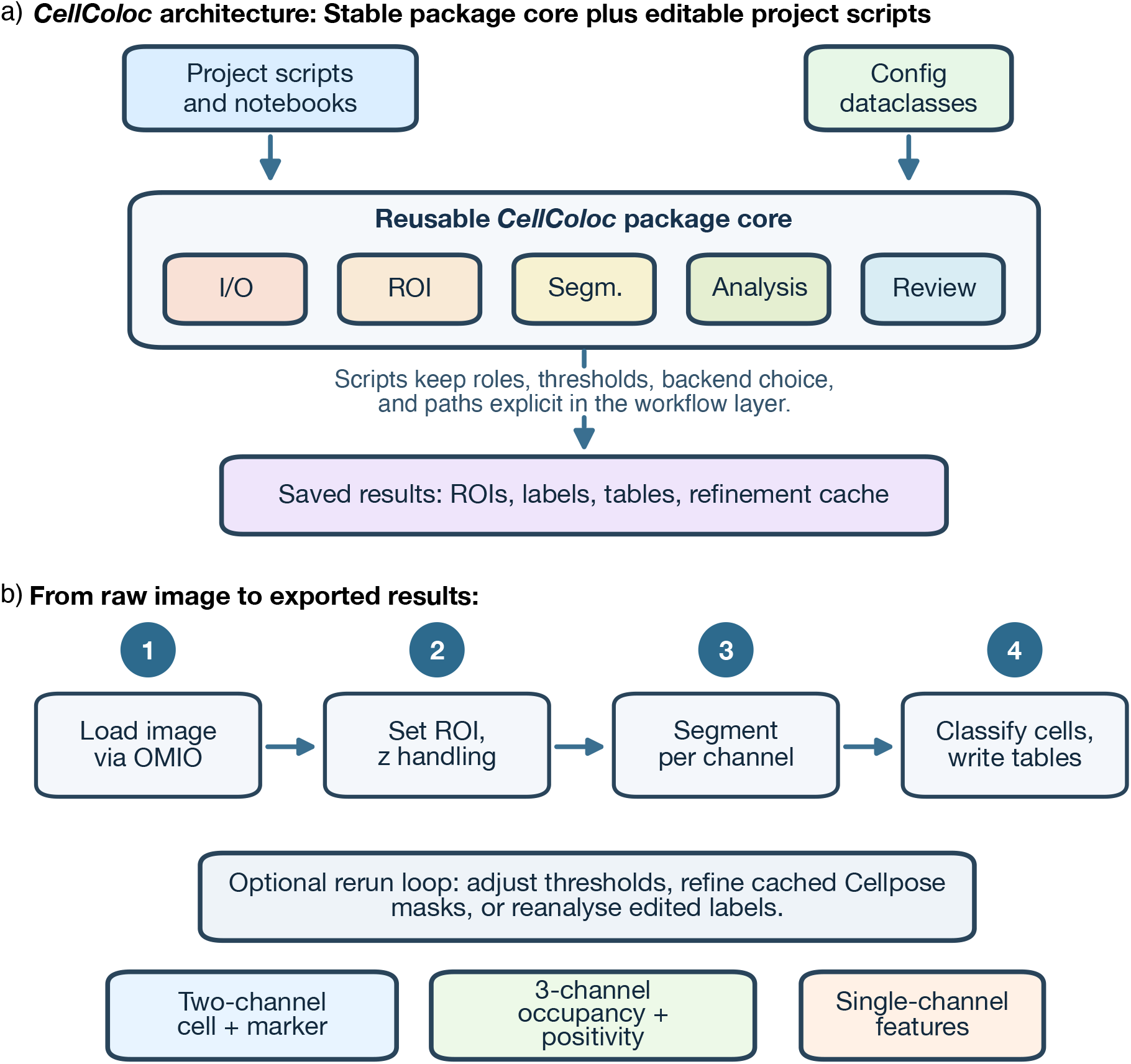
CellColoc architecture and workflow model. (a) The package follows an explicit script-driven progression from OMIO-based loading through ROI definition, channel-wise segmentation, overlap quantification, optional refinement, and standardized export. (b) Reusable project-specific scripts remain separate from stable package modules for I/O, ROI handling, segmentation, analysis, visualization, and export. Reproducibility further relies on persistent intermediate artifacts, including ROI masks, per-channel label images, positive-cell masks, and structured tabular summaries.

Figure 1 a) emphasizes the operational sequence: microscopy data are loaded, a region of interest is defined, segmentation is performed channel by channel, and overlap calls and summary tables are written out together with the intermediate masks needed for review or reruns. This makes the workflow itself inspectable at the same level at which users actually adapt it.

Figure 1 b) shows the typical end-user workflow arrangement in which a small user script imports the package, defines paths and parameters, and calls the main analysis function. The user-workflow is deliberately kept separate from the package logic shown in Figure 1 a), which is versioned and maintained independently. In particular, configuration is expressed through plain dataclasses, including separate objects for channel assignment, segmentation settings, overlap thresholds, display names, and runtime behavior. This keeps project-specific choices explicit and editable while preserving a stable reusable analysis core. Design Goal 4 in Box 1 is implemented through OMIO, which allows the same scripts to operate on formats such as TIFF, OME-TIFF, Zeiss CZI, Zeiss LSM or even Thorlabs raw files while preserving metadata access and policy-driven microscopy I/O behavior [7, 8]. Each configured analysis channel can independently choose its segmentation backend. Currently supported backends are Cellpose, Otsu thresholding, Li thresholding, and percentile-based thresholding, while CellColoc’s modular design allows new backends to be added without changing the overall workflow. Optional prefilters and postfilters can be applied per channel, and analysis preparation can include z-cropping or z-projection when a 2D view of volumetric data is desired. ROIs can be drawn interactively in napari [15], reused from previously saved ROI masks, or instantiated as full-image regions. For volumetric data, CellColoc currently uses 2D ROI masks that are broadcast along the z axis. This keeps region definition inspectable in routine workflows while still enabling 3D segmentation and quantification within the selected subvolume.

### Two-channel object-based colocalization

The core CellColoc procedure is an object-based positivity call, i.e., one segmented channel defines the reference cell objects, another segmented channel defines the marker support, and positivity is assigned from their overlap under explicit thresholds. Let a microscopy stack be represented as

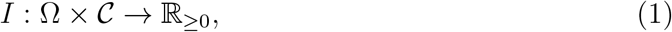

where Ω ⊆ ℤ^*d*^ is the spatial lattice, *d* ∈ *{*2, 3*}* is the effective dimensionality after preprocessing, and *C* indexes image channels. Optional z-cropping or z-projection is represented by a preprocessing operator *P* , yielding analysis images

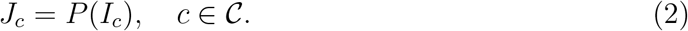

For one region of interest *r*, let *R*_*r*_ : Ω → *{*0, 1*}* denote the ROI indicator and Ω_*r*_ = *{x* ∈ Ω : *R*_*r*_(*x*) = 1*}* the selected analysis domain. For each configured channel *c*, CellColoc applies a channel-specific segmentation operator

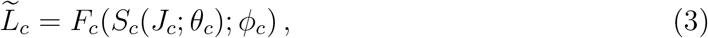

where *S*_*c*_ is the chosen backend with parameters *θ*_*c*_ and *F*_*c*_ denotes optional postprocessing with parameters *ϕ*_*c*_.

In the two-channel mode, the primary cell channel *c*_cell_ defines cell objects

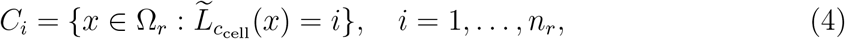

whereas the marker channel *c*_mark_ defines a marker-support set

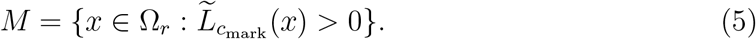

CellColoc interprets colocalization as an object-based overlap decision. For each cell object it computes the absolute overlap

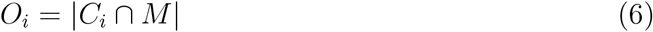

and the within-cell overlap fraction

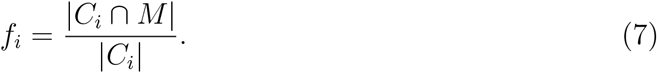

With absolute and fractional thresholds *τ*_abs_ and *τ*_frac_, the marker-positivity call becomes

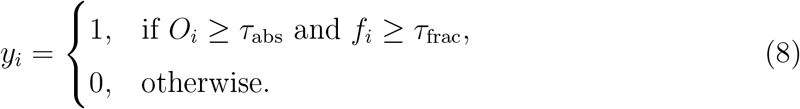

ROI-level summaries then follow naturally, for example

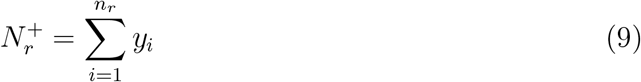

for the number of marker-positive cells and

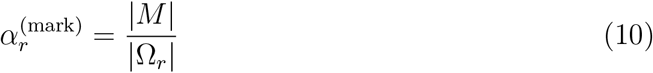

for marker occupancy within the ROI. Because the segmentation backend appears only inside *S*_*c*_, learned segmentation on one channel can be combined with threshold-based segmentation on another without changing the downstream decision rule.

### Three-channel occupancy and extended positivity

The three-channel mode extends the same formalism by introducing an additional segmented support set

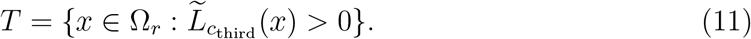

At ROI level, this channel contributes occupancy summaries such as

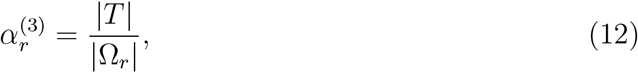

with corresponding 3D volume fractions when the analysis remains volumetric. At the per-cell level, CellColoc can compute a third-channel occupancy fraction

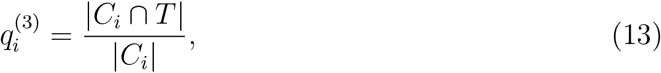

and, if enabled for a given workflow, a third-channel positivity call

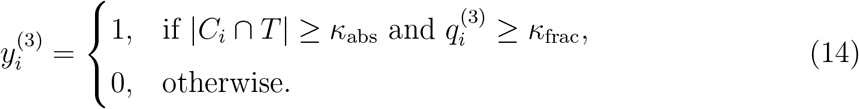

Double-positive states then follow from

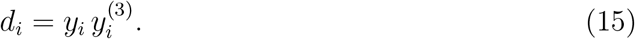

This mode is useful when an additional channel represents an infiltrating, territorial, or lesion-associated structure whose occupancy within the ROI is scientifically relevant and may also be interpreted at the cell level.

### Single-channel segmentation and feature extraction

Not every microscopy project requires a cross-channel positivity call, but the same segmentation, ROI, and export machinery remains useful for counting and morphology analysis. Let

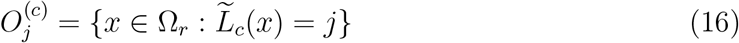

denote one segmented object from an arbitrary channel *c*. For every segmented object, CellColoc exports a position, morphology, and intensity vector

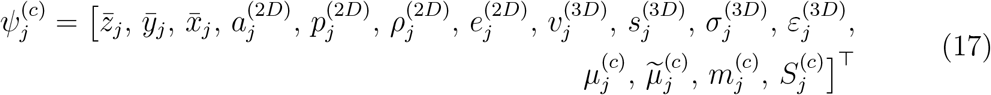

with entries populated according to the effective dimensionality. In effective 2D analyses, roundness is reported as

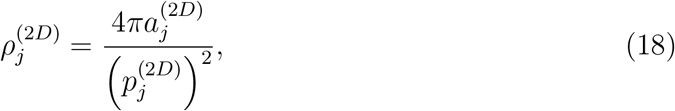

where 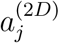 and 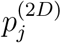 are object area and perimeter. In 3D, sphericity is reported as

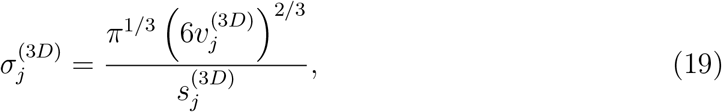

where 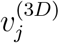 and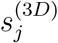 are object volume and surface area. Intensity readouts are measured in the native intensity units of the analyzed channel image: 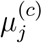 is mean intensity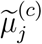 median intensity, 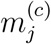 maximum intensity, and 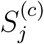 integrated intensity, i.e., the sum of voxel or pixel values inside the segmented object.

This feature export model is not limited to the dedicated single-channel branch. In multi-channel analyses, CellColoc writes corresponding object-wise property tables for every segmented channel, including the primary cell channel, the marker channel, and, when present, the optional third channel. The dedicated single-channel mode simply omits the positivity variables 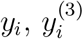, and *d*_*i*_ while reusing the same segmentation and measurement schema.

### Persistent outputs and rerun semantics

Design Goal 3 in Box 1 is implemented by treating export as part of the workflow model rather than as a final afterthought. For each ROI, the package writes a structured output collection

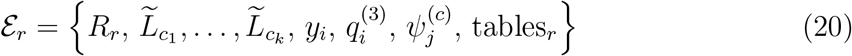

to a standardized `results/` directory next to the source dataset. Multi-channel analyses export detailed overlap tables, per-cell summaries, and ROI-level overviews; single-channel analyses export per-object morphology and intensity tables and ROI summaries. Importantly, intermediate masks and cached Cellpose outputs can be revisited for threshold refinement, and edited labels can be reanalyzed without rebuilding the workflow from scratch. The main exported artifact families are summarized in Table 2. These persistent intermediates are central to the package’s transparency and reuse model.

**Table 2.**
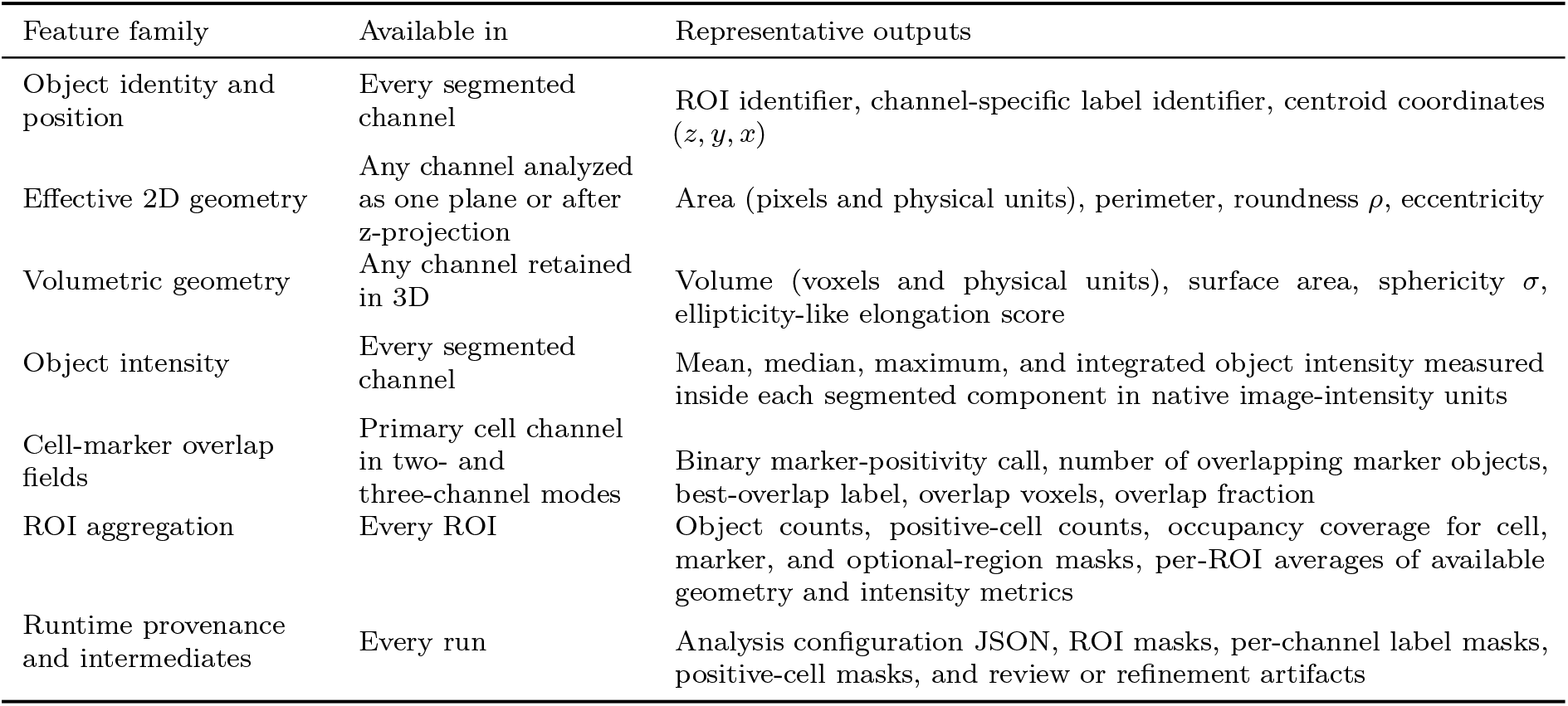
Representative CellColoc exports and feature families. The same object-level geometry and intensity schema is reused for every segmented channel; the multi-channel cell table additionally stores overlap-specific classification fields.

| Feature family | Available in | Representative outputs |
| --- | --- | --- |
| Object identity and position | Every segmented channel | ROI identifier, channel-specific label identifier, centroid coordinates $(z, y, x)$ |
| Effective 2D geometry | Any channel analyzed as one plane or after z-projection | Area (pixels and physical units), perimeter, roundness $\rho$, eccentricity |
| Volumetric geometry | Any channel retained in 3D | Volume (voxels and physical units), surface area, sphericity $\sigma$, ellipticity-like elongation score |
| Object intensity | Every segmented channel | Mean, median, maximum, and integrated object intensity measured inside each segmented component in native image-intensity units |
| Cell-marker overlap fields | Primary cell channel in two- and three-channel modes | Binary marker-positivity call, number of overlapping marker objects, best-overlap label, overlap voxels, overlap fraction |
| ROI aggregation | Every ROI | Object counts, positive-cell counts, occupancy coverage for cell, marker, and optional-region masks, per-ROI averages of available geometry and intensity metrics |
| Runtime provenance and intermediates | Every run | Analysis configuration JSON, ROI masks, per-channel label masks, positive-cell masks, and review or refinement artifacts |

### Synthetic benchmark generation and evaluation metrics

To evaluate CellColoc under controlled conditions, we generated a synthetic three-channel benchmark with stored ground truth for object counts, colocalization calls, morphology, and area coverage. The benchmark was designed to exercise the same workflow elements used in ordinary analyses: channel-wise segmentation, object-based positivity calling, optional third-channel occupancy measurement, and export of per-object and per-stack tables. Two renderings were generated from the same underlying object layout. In the first, channels 0 and 1 were rendered as sharply bounded filled objects; in the second, the same supports were rendered with softer Gaussian-intensity profiles before noise was added. The sharp rendering tests performance when object boundaries are unambiguous, whereas the Gaussian rendering probes robustness when boundaries become less well defined.

For each synthetic stack, the number of channel-0 objects was sampled uniformly from 10 to 30. Each ground-truth object was defined as a rotated filled support

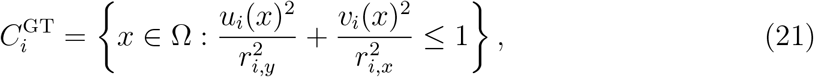

where *x* ∈ Ω ⊂ ℤ^2^, (*u*_*i*_, *v*_*i*_) are coordinates in the object’s rotated frame, and the effective diameter range was constrained to 15–35 pixels by the sampled semi-axes. In the sharp rendering, channel 0 was generated as

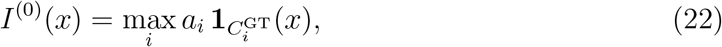

with amplitudes *a*_*i*_ and object centres sampled so that the supports did not merge under the benchmark placement rule.

Channel 1 was generated from the same number of filled objects. A fraction of approximately 70% reused channel-0 object locations as seeds and received small random offsets, so that their support sets intersected the corresponding channel-0 objects. The remaining approximately 30% were placed away from all channel-0 supports and already-placed marker objects. For the softer-boundary variant, the same object supports were instead rendered with Gaussian intensity falloff

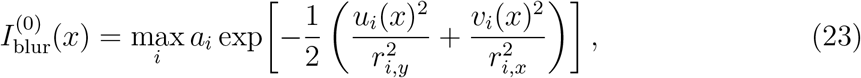

and analogously for channel 1. Channel 2 was intentionally not object-based. Instead, it was derived from a smoothed Gaussian random field *h*(*x*) whose upper level set

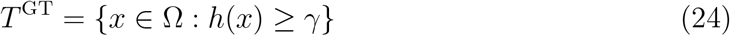

was thresholded and morphologically cleaned to occupy approximately 30% of the field of view. The final benchmark images were obtained after additive noise and per-channel intensity scaling, while the ground-truth masks and tables were stored separately for both renderings. Because this third channel represents a contiguous area-like structure rather than discrete cells, its occupancy was evaluated with threshold-based segmentation rather than Cellpose-based object parsing.

Ground-truth positivity in the benchmark followed the same object-based logic used by CellColoc. For every ground-truth channel-0 object, the best-overlap channel-1 label was identified, and the cell was counted as marker-positive if both

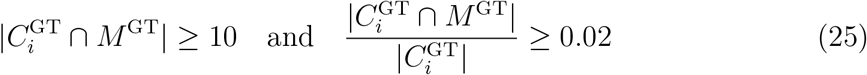

held. Ground-truth summary tables further recorded image area, per-channel object counts, the number of marker-positive channel-0 objects, channel-2 coverage, and per-object morphology including area, eccentricity, and roundness.

Benchmark evaluation compared CellColoc outputs against this stored ground truth on both per-stack and matched-instance levels. For the sharply bounded rendering, we evaluated both a threshold-based branch and a Cellpose-based branch on the two object channels; the Gaussian rendering was used to assess how softer boundaries influence detection and morphology estimates. Instance matching between ground-truth labels *G*_*u*_ and predicted labels *P*_*v*_ used the IoU matrix

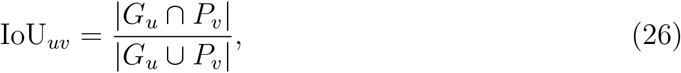

followed by one-to-one assignment and acceptance only for pairs with IoU_*uv*_ ≥ 0.5. From the resulting numbers of true positives (*TP*), false positives (*FP*), and false negatives (*FN*), we computed

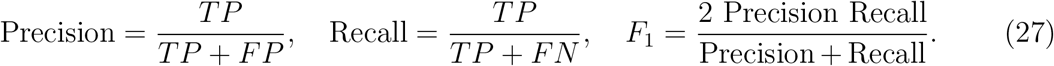

At the per-stack level, counts and occupancy were summarized through absolute errors

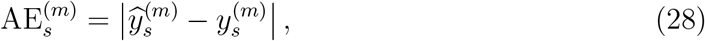

where *m* indexed channel-0 counts, channel-1 counts, marker-positive channel-0 counts, and channel-2 coverage, respectively. Mean absolute errors were then

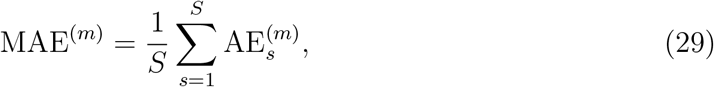

with coverage errors reported in percentage points. For one-to-one matched channel-0 objects, we additionally reported morphology agreement using the mean absolute error, median absolute error, and Pearson correlation between ground-truth and predicted object areas and roundness values.

## Results

### Synthetic benchmark with known ground truth

To test whether CellColoc can recover scientifically relevant counts, overlap calls, and morphology summaries from a reproducible scripted workflow, we generated a synthetic benchmark of 20 three-channel OME-TIFF stacks. Channel 0 contained 10–30 noisy, fully filled cell-like blobs with variable diameters (15–35 pixels), channel 1 reused the first-channel locations such that approximately 70% of objects overlapped channel-0 cells and 30% were placed away from them, and channel 2 contained an irregular area occupying roughly 30% of the image. Ground-truth tables stored per-stack object counts, channel-1-positive channel-0 counts, object areas, roundness values, eccentricities, and channel-2 coverage. The same script-driven CellColoc workflow used for ordinary analyses was then applied to all stacks, and instance-level comparisons used one-to-one IoU matching with a threshold of 0.5 for the benchmark-specific evaluation.

Representative synthetic inputs and the corresponding CellColoc-derived masks are shown in Figure 2 a–h. In this primary benchmark, channels 0 and 1 were segmented with default `cpsam`-based Cellpose settings, whereas the contiguous channel-2 region was segmented by Otsu thresholding because it represents an area-like support rather than a population of discrete cells. Under this mixed-backend configuration, CellColoc recovered channel-0 counts perfectly across all 20 stacks (mean absolute error, 0.00 cells per stack) and recovered channel-1 counts with a mean absolute error of only 0.20 cells per stack. The number of channel-1-positive channel-0 cells was recovered with a mean absolute error of 0.05 cells per stack, and channel-2 occupancy was estimated with a mean absolute error of 1.13 percentage points.

**Figure 2:**
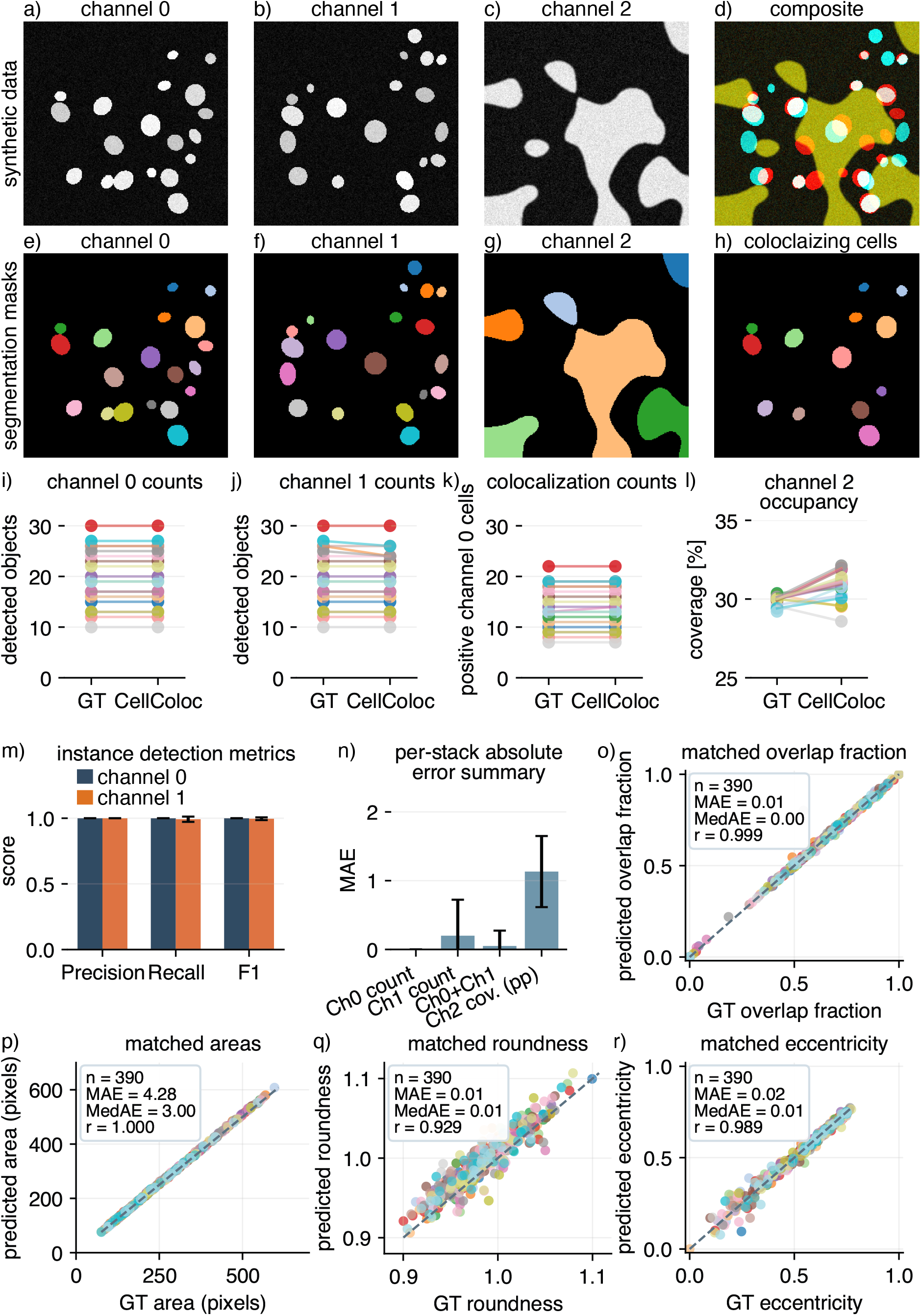
Cellpose-based synthetic benchmark on sharp filled-object stacks. Channels 0 and 1 were segmented with default `cpsam`-based Cellpose settings, whereas the non-cellular channel-2 region was segmented by Otsu thresholding. (a–d) Representative synthetic benchmark stack shown as channel 0, channel 1, channel 2, and a composite view. (e–h) Corresponding CellColoc masks for the cell, marker, region, and positive-cell outputs. (i–l) Ground truth versus CellColoc estimates for channel-0 counts, channel-1 counts, channel-1-positive channel-0 counts, and channel-2 occupancy. Each line connects the same stack across the two summaries. (m) Mean instance-detection precision, recall, and F1 scores for channels 0 and 1. (n) Mean absolute error summaries across the four benchmark targets. (o) One-to-one matched channel-0 overlap fractions. (p) One-to-one matched channel-0 object areas. (q) One-to-one matched channel-0 roundness values. (r) One-to-one matched channel-0 eccentricity values. Point colors encode stacks.

Instance-level agreement was correspondingly strong (Figure 2 i–n). For channel 0, mean precision, recall, and F1 were all 1.000. For channel 1, mean precision remained 1.000, whereas recall and F1 were 0.992 and 0.996, respectively. The residual channel-1 count error arose from only four missed marker objects across three stacks. Two of these unmatched objects were among the smallest in the benchmark (72–88 pixels), and all four occurred in locally crowded marker configurations in which nearby supports lay within roughly 10–16 pixels of the missed object centroid. A likely explanation is therefore that Cellpose occasionally merges a small or closely apposed marker object into a neighbor rather than hallucinating extra detections. Importantly, this residual marker undersegmentation had only a minimal effect on the downstream colocalization readout, as reflected by the 0.05-cell mean absolute error for marker-positive channel-0 cells.

Matched overlap and morphology readouts were likewise preserved at high fidelity (Figure 2 o–r). Across 390 one-to-one matched channel-0 objects, predicted overlap fractions closely tracked the stored ground truth, with a mean absolute error of 0.0054, a median absolute error of 0.0027, and a Pearson correlation of *r* = 0.999. Predicted and ground-truth areas showed a mean absolute error of 4.28 pixels, a median absolute error of 3 pixels, and an almost perfect Pearson correlation of *r* = 0.9998. Roundness estimates remained close, with a mean absolute error of 0.0140, a median absolute error of 0.0130, and a Pearson correlation of *r* = 0.929, while eccentricity estimates yielded a mean absolute error of 0.0152, a median absolute error of 0.0097, and a Pearson correlation of *r* = 0.989. Taken together, these results show that the sharp filled-object benchmark is recovered not only at the count level but also at the level of per-object overlap and morphology.

The supplementary benchmarks help interpret where performance begins to soften. Supplementary Figure S2 shows the same sharp filled-object stacks analyzed with threshold-based segmentation on the two object channels. That variant still recovered counts and colocalization at a useful level, but its morphology agreement was weaker than in the Cellpose-based benchmark. Supplementary Figure S1 then moves one step further by rendering the same object layout with softer Gaussian boundaries before noise was added. Under these less sharply defined conditions, CellColoc still delivered strong cell and marker counts, but the matched overlap and shape descriptors became more sensitive to contour placement: areas were systematically underestimated, and the roundness and eccentricity summaries shifted modestly as boundaries became less sharply defined. This is consistent with the fact that both thresholding and learned object models must place a discrete contour on an increasingly ambiguous intensity boundary. The main practical point is that cell counts and positivity calls remain highly stable, whereas fine per-object boundary-derived readouts are the first quantities to drift as object edges become less well defined.

### Biological example: regional microglia analysis in CA1 and cortex

To test the workflow on real microscopy data, we analyzed a multi-stack Zeiss CZI dataset of tdTomato-labeled microglia with accompanying Iba1 immunostaining and DAPI counterstaining. In this dataset, tdTomato marks the recombined CX3CR1-positive microglial population, Iba1 provides an independent microglia/macrophage-lineage marker, and DAPI serves mainly for anatomical orientation and quality control. We focused the comparison on two anatomical groups, hippocampal CA1 and cortical CTX stacks as indicated in Figure 3 b) (created with anatomical guidance from public mouse brain atlas references [16, 17] and used only to indicate the approximate CTX and CA1 acquisition fields), and used CellColoc in its two-channel configuration to ask how consistently tdTomato-positive cells were also Iba1-positive across these two regions.

**Figure 3:**
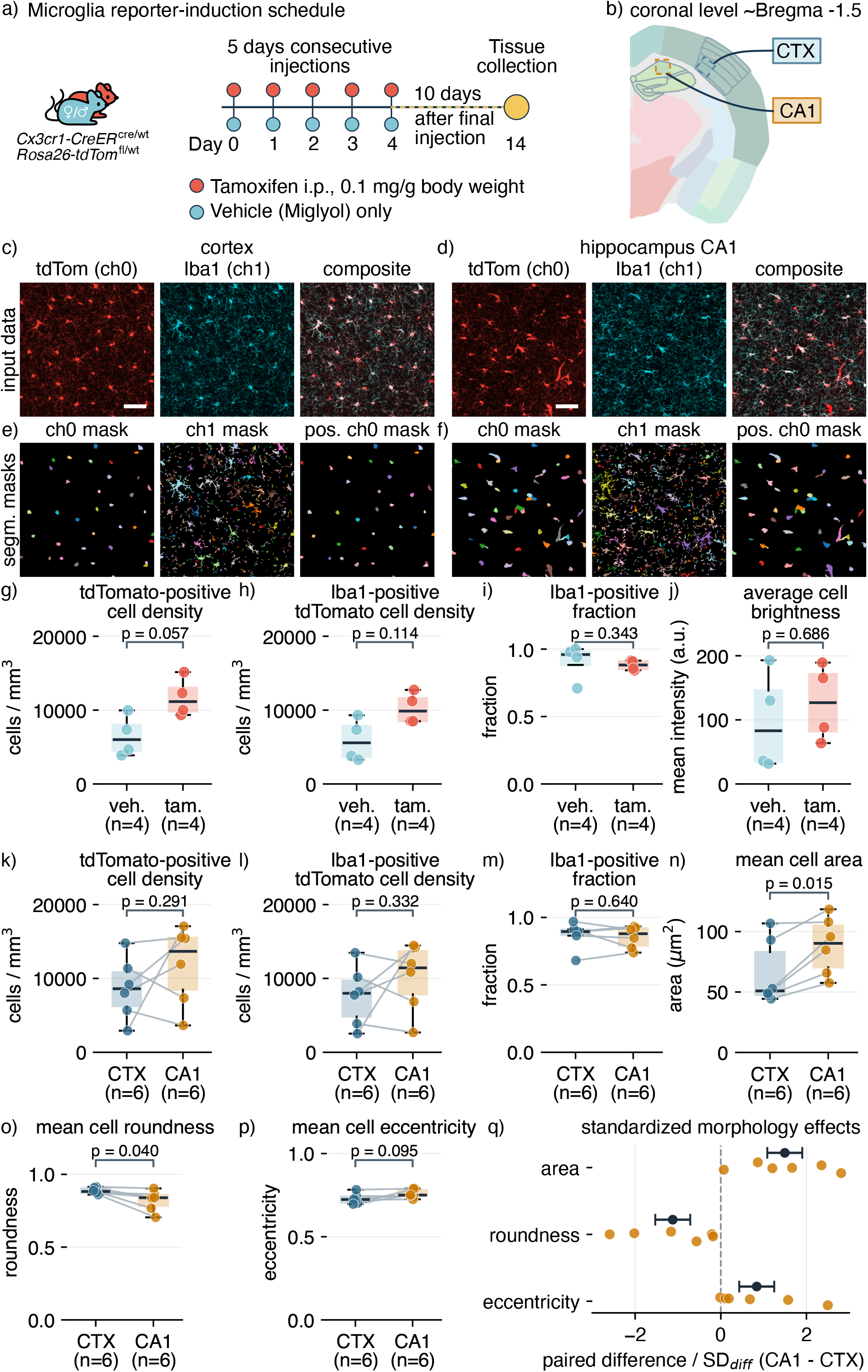
Biological microglia batch-analysis example. (a) Experimental schedule showing microglia reporter-induction vehicle or tamoxifen administration and tissue collection time points. (b) Brain-section schematic with sampling boxes indicating approximate cortical and hippocampal CA1 imaging fields (drawn by the authors with anatomical guidance from public mouse brain atlas references [16, 17]). (c) Three representative cortex panels showing, from left to right, tdTomato-labeled microglia cells (channel 0), Iba1 staining of the microglia (channel 1), and the composite cortex view. (d) Analogous hippocampal CA1 view. (e) Three cortex result panels showing, from left to right, the channel-0 Cellpose segmentation mask, the channel-1 Otsu segmentation/thresholding mask, and the channel-1-positive channel-0 mask. (f) Analogous CA1 result view. (g–j) Treatment-sensitivity analysis: (g) Mouse-level tdTomato-positive cell density in vehicle- and tamoxifen-treated animals. (h) Mouse-level Iba1-positive tdTomato cell density in vehicle- and tamoxifen-treated animals. (i) Mouse-level Iba1-positive fraction in vehicle- and tamoxifen-treated animals. (j) Mouse-level average tdTomato cell brightness in vehicle- and tamoxifen-treated animals. For panels g–j, repeated scans were first averaged within mouse and region; the available regional values were then averaged per mouse to obtain one treatment-level value per animal. This yielded four vehicle-treated and four tamoxifen-treated mice (*n* = 4 vs. *n* = 4); vehicle-only and tamoxifen-treated animals were compared with two-sided Mann–Whitney U tests. Tamoxifen-treated mice showed a trend toward higher tdTomato-positive cell density (*p* = 0.057) and a weaker non-significant tendency toward higher Iba1-positive tdTomato cell density (*p* = 0.114), whereas the Iba1-positive fraction and average tdTomato cell brightness did not differ detectably between treatment groups (*p* = 0.343 and *p* = 0.686, respectively). (k–q) Regional comparison of CTX and CA1 microglia readouts with treatment groups pooled: (k) tdTomato-positive cell density for the regional CTX/CA1 comparison. (l) Iba1-positive tdTomato cell density (Iba1 microglia colocalization). (m) Iba1-positive fraction. (n) Mean cell area. (o) Mean cell roundness. (p) Mean cell eccentricity. (q) Standardized paired morphology effects, computed as CA1–CTX paired differences divided by the standard deviation of the paired differences for area, roundness, and eccentricity. The paired regional comparison comprised six C57BL/6J *Cx3cr1-creER*^cre/wt^; *Rosa26_tdTom*^fl/wt^ mice pooled across sexes (3 male, 3 female; mean age 40.5 weeks), because only mice with matched CTX and CA1 acquisitions could contribute to the paired regional test (see Supplementary Table S1). Cell densities are normalized to the imaged stack volume and reported in cells/mm^3^. Repeated scans from the same mouse and region were averaged before testing. Normality of paired CA1–CTX differences was assessed per endpoint; normally distributed endpoints were tested with paired *t*-tests and the remaining endpoints with Wilcoxon signed-rank tests. Additional cohort and acquisition details are provided in the Supplementary material. Scalebar: 50 *µ*m.

Each CZI stack was loaded via OMIO and max-z projected to a 2D analysis plane. For the CA1 stacks, the tdTomato channel was lightly prefiltered with a Gaussian filter (*σ*_*xy*_ = 0.5) before Cellpose segmentation to emphasize soma-scale signal in the projection and attenuate fine processes; the Iba1 channel was analyzed with threshold-based marker masking. The resulting masks, tables, and preview panels were then exported with the same workflow used elsewhere in CellColoc.

Because the biological example included both tamoxifen- and vehicle-treated mice (Figure 3 a)), we first asked whether treatment introduced an obvious shift in the key tdTomato/Iba1 readouts that would preclude pooling animals for the regional comparison. In this mouse-level sensitivity analysis, repeated scans were averaged within region and the available regional values were then averaged per mouse to obtain one treatment-level value per animal. Figure 3 g)–j) shows a trend toward higher tdTomato-positive cell density after tamoxifen treatment (*p* = 0.057) and a similar but weaker non-significant trend for Iba1-positive tdTomato cell density (*p* = 0.114), whereas the Iba1-positive fraction remained high and did not differ detectably between treatment groups (*p* = 0.343). Average tdTomato cell brightness likewise showed no statistically detectable treatment-associated shift in this small cohort (*p* = 0.686). We therefore analyzed the regional CellColoc example with treatment groups pooled.

For regional comparisons, tdTomato-positive cell counts and Iba1-positive tdTomato-positive cell counts were normalized by the imaged stack volume and reported as cell densities in cells/mm^3^. Figure 3 c) and d) show representative CTX and CA1 input views for tdTomato, Iba1, and the corresponding overlay, whereas Figure 3 e) and f) show the derived channel-0, channel-1, and positive-cell masks for the same representative stacks. Figure 3 k)–q) summarizes the corresponding region-level quantitative readouts. Because no ground-truth labels are available for these biological data, the purpose of this example is not absolute benchmarking but to demonstrate the intended use of CellColoc on experimental microscopy data.

The quantitative regional comparisons in Figure 3 k)–m) show that most tdTomato-positive cells were also classified as Iba1-positive in both regions. The mean Iba1-positive fraction remained high in both regions (0.869 in CTX and 0.854 in CA1). Volume-normalized tdTomato-positive cell density and Iba1-positive tdTomato-positive cell density were numerically higher in CA1 than in CTX, but neither difference was statistically significant (*p* = 0.291 and *p* = 0.332, respectively), and the positivity fraction itself likewise remained indistinguishable between regions (*p* = 0.640). In contrast, the morphology readouts shown in Figure 3 n)–q) separated the regions more clearly. CA1 microglia showed a larger mean projected cell area than CTX microglia (88.4 versus 65.6 *µ*m^2^, *p* = 0.015) together with lower roundness (0.819 versus 0.887, *p* = 0.040), indicating broader and less circular projected cell profiles in CA1. Mean cell eccentricity showed the corresponding tendency toward more elongated CA1 profiles (0.758 versus 0.729, *p* = 0.095), but this trend did not cross the significance threshold in the present cohort. The standardized paired morphology summary in Figure 3 q) combines these three endpoints on a common effect scale and illustrates the shared direction of the regional morphology shift: CA1 cells were larger and less round, with a matching tendency toward higher eccentricity. Taken together, these results show that CellColoc yields biologically interpretable per-cell and per-region readouts even on structurally complex cell types such as microglia and can support region-wise comparisons of marker positivity, density, and morphology within a single transparent analysis workflow.

## Discussion

The central contribution of CellColoc is that it turns segmentation-based microscopy analyses into a stable, inspectable workflow pattern rather than a collection of one-off scripts. The controlled synthetic benchmark shows that this workflow can deliver quantitatively reliable outputs when ground truth is known. In the Cellpose-based sharp-object benchmark, channel-0 counts were recovered exactly across all stacks, channel-1 counts and colocalization calls were nearly exact, and object morphology remained tightly aligned with ground truth. These results are important not only because the underlying segmentation is strong, but because the full workflow surrounding it remains transparent: the same run writes the masks, positivity assignments, ROI summaries, and object-level tables that a user would later inspect, archive, or reanalyze.

The biological microglia example extends this point from controlled data to a real application. After establishing in the synthetic benchmark that CellColoc can recover counts, overlap calls, and morphology with high fidelity, the tdTomato/Iba1 dataset shows that the same workflow can be used to obtain biologically interpretable measurements from experimental images.

The treatment-sensitivity analysis provides useful context for interpreting the tdTomato reporter channel before pooling animals for the regional comparison. Tamoxifen activates CreER-mediated recombination rather than simply increasing fluorescence of already recombined cells, and substantial tamoxifen-independent recombination has been reported for some microglial CreER reporter combinations, with recombination efficiency and spontaneous reporter labeling depending on the specific CreER line and reporter allele [18, 19]. In our cohort, tamoxifen-treated animals showed a trend toward higher tdTomato-positive cell density and a similar non-significant tendency for Iba1-positive tdTomato cell density, consistent with a possible increase in reporter-labeling extent or detectability. By contrast, the fraction of tdTomato-positive objects classified as Iba1-positive remained high and statistically indistinguishable between vehicle- and tamoxifen-treated mice, and average tdTomato cell brightness also did not differ detectably. This pattern does not establish treatment equivalence, but it indicates that the observed tdTomato-positive population retained a similar microglial marker correspondence across treatment groups, without an accompanying treatment-associated shift in per-cell reporter brightness in this dataset.

For the paired CTX-versus-CA1 comparison, CellColoc reports Iba1 positivity among tdTomato-positive cells, compares volume-normalized regional cell densities, and quantifies morphology differences on a mouse-matched basis. Within this cohort, the regional pattern was driven more strongly by morphology than by density or marker-positivity frequency: the volume-normalized tdTomato and Iba1-positive tdTomato densities showed only non-significant CA1-increasing trends, whereas CA1 cells had significantly larger projected areas and significantly lower roundness, together with a matching trend toward higher eccentricity. These observed regional differences in morphology are consistent with previous reports of microglial heterogeneity in distribution and morphology across the adult brain [20], with later work showing that region-specific microglial phenotypes are maintained by local cues [21] and may be especially diverse in hippocampal contexts [22]. The comparatively modest regional differences in cell density likewise agree with reports that microglial densities can remain similar across brain structures despite differences in other phenotypic features [23]. Even without ground-truth labels, this is therefore a meaningful demonstration of scientific utility: CellColoc enables known and biologically plausible microglial readouts to be reproduced quickly from experimental microscopy data while preserving reusable cell-level and region-level outputs for inspection, reanalysis, and downstream reporting.

The additional variants of the synthetic benchmark further show how performance changes under less favorable segmentation conditions. When object boundaries are crisp but threshold-based rather than Cellpose-based segmentation is used, counts remain useful but morphology agreement becomes weaker. When the same objects are rendered with softer Gaussian boundaries, cell and marker counts still remain strong, yet area tends to be underestimated and roundness mildly overestimated. This pattern is expected since, as edges become less well defined, the exact contour placement becomes more sensitive to threshold choice or to how a learned model resolves diffuse local contrast. In practical terms, CellColoc therefore appears most robust for the readouts that many users care about first, namely cell counts, marker positivity, and occupancy, whereas fine-grained boundary-derived morphology is the quantity most likely to drift when segmentation conditions become ambiguous. These observations also define the package’s current boundaries. Biological interpretation still depends on choosing an appropriate segmentation backend, preprocessing strategy, and overlap threshold for the dataset at hand.

The present results show that CellColoc provides a strong and useful workflow core for microscopy projects that require segmentation-based cell quantification. Its main practical advantages are the combination of flexible segmentation-backend choice, transparent per-cell overlap classification, persistent intermediate outputs, explicit rerun semantics, and rapid export of cell counts, colocalization calls, occupancy summaries, and morphology tables. In practice, this replaces ad hoc manual counting and opaque spreadsheet workflows with an analysis structure that is scriptable, inspectable, and reproducible from raw image to final table. Just as importantly, the workflow surface remains small enough to be extended without destabilizing the package: new segmentation backends, dataset-specific batch scripts, postfilters, or richer exports can be added while preserving the same explicit analysis model. This makes CellColoc a practical and extensible foundation for transparent, reproducible, and scientifically useful cell colocalization workflows in microscopy research.

## Data and code availability

CellColoc is distributed under the GNU General Public License v3.0 or later. The archived software release is publicly available via Zenodo [24], and the source code, documentation, and issue tracker are hosted in the public repository (https://github.com/fabriziomusacchio/CellColoc). The accompanying example-data archive [25] includes both the synthetic benchmark and the microglia dataset, together with the corresponding ground-truth tables, CellColoc masks, and exported summary tables. The repository additionally contains a full set of example scripts that reproduce the analyses shown in this paper, including the synthetic benchmark, the microglia batch analysis, and the central figure-generation code.

## Acknowledgments

We thank the DZNE Animal Research Facility (ARF) for animal care and husbandry, and the DZNE Light Microscopy Facility (LMF) for support with confocal imaging. Both ARF and LMF facilitated the biological microglia experiment and provided technical support for the acquisition of the dataset.

## Contributions

Fabrizio Musacchio developed and benchmarked the CellColoc pipeline, wrote tests, tutorials, and documentation, and wrote the manuscript. Sophie Crux planned the microglia experiment and performed the tamoxifen injections. Denise Maria Hoffmann and Felix Nebeling performed the perfusions. Henrike Antony and Arush Baijal carried out staining, mounting, and confocal imaging. Martin Fuhrmann supervised the study.

## Supplementary figures

**Figure S1:**
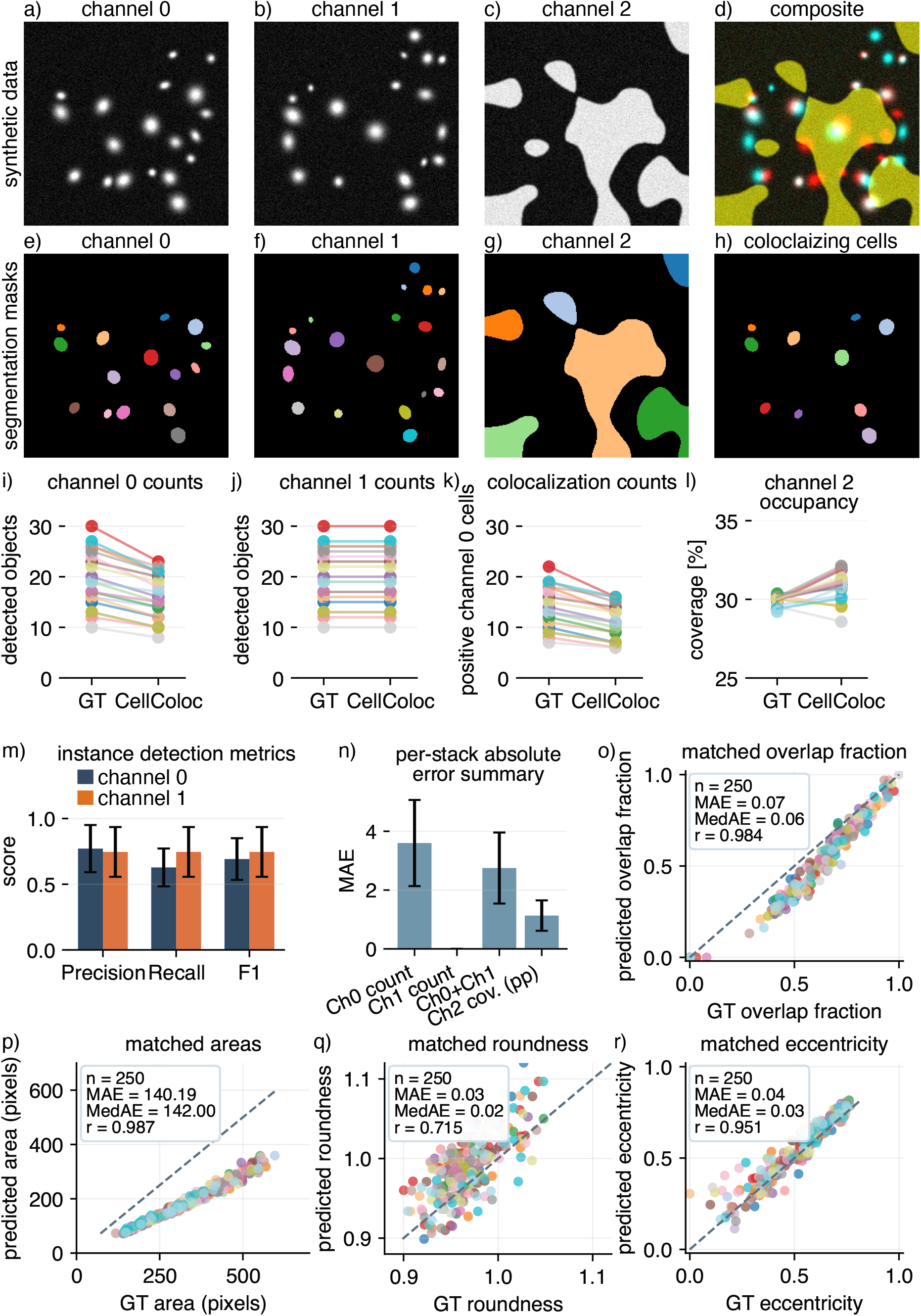
Gaussian-intensity synthetic benchmark variant. The underlying support geometry matches the main benchmark on the synthetic data shown in Figure 2, but channels 0 and 1 were rendered with softer Gaussian intensity falloff before additive noise and Otsu segmentation. (a–d) Representative benchmark stack shown as channel 0, channel 1, channel 2, and a composite view. (e–h) Corresponding CellColoc masks for the cell, marker, region, and positive-cell outputs. (i–l) Ground truth versus CellColoc estimates for channel-0 counts, channel-1 counts, channel-1-positive channel-0 counts, and channel-2 occupancy. Each line connects the same stack across the two summaries. (m) Mean instance-detection precision, recall, and F1 scores for channels 0 and 1. (n) Mean absolute error summaries across the four benchmark targets. (o) One-to-one matched channel-0 overlap fractions. (p) One-to-one matched channel-0 object areas. (q) One-to-one matched channel-0 roundness values. (r) One-to-one matched channel-0 eccentricity values. Point colors encode stacks.

**Figure S2:**
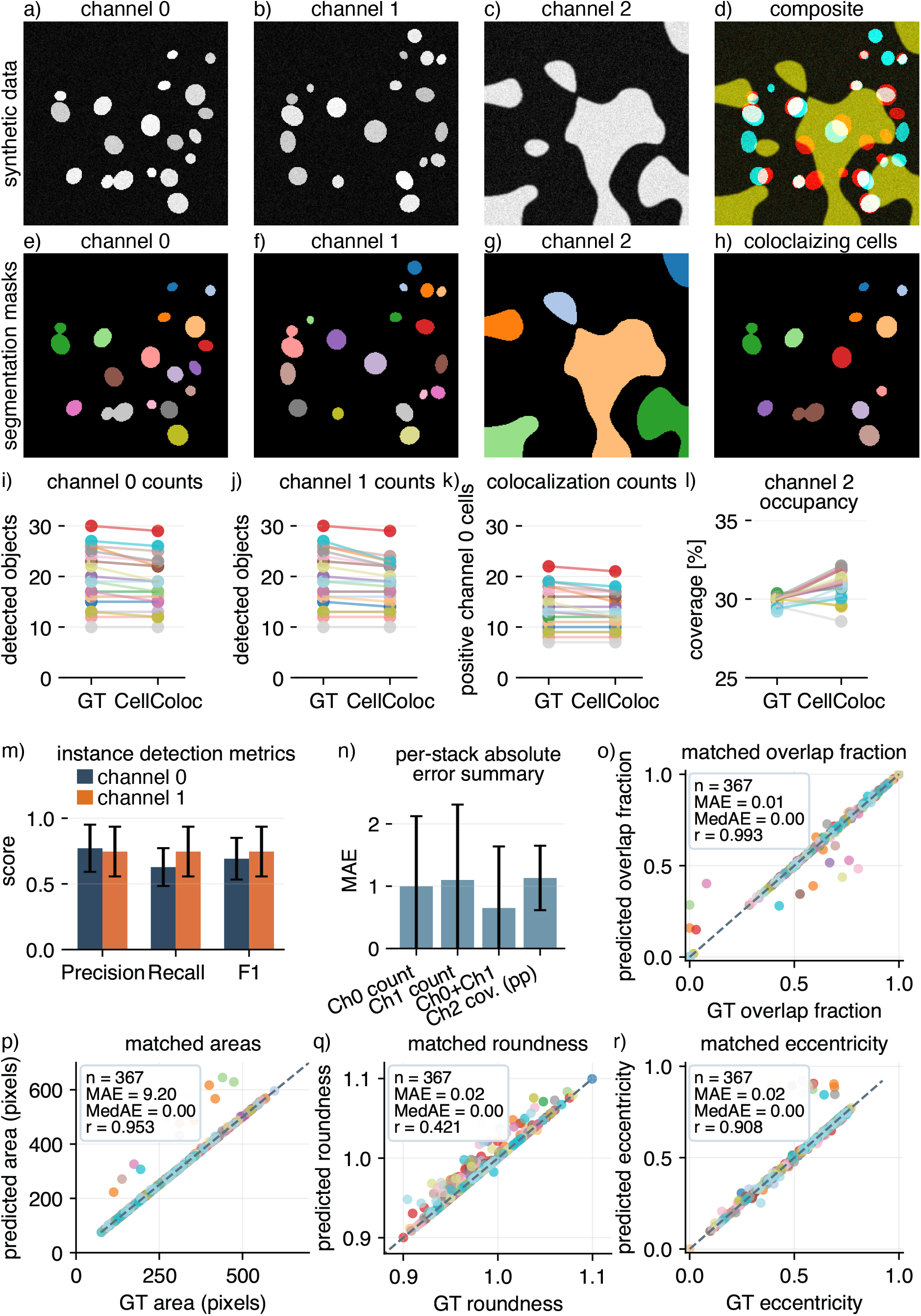
Threshold-based synthetic benchmark variant on the sharp filled-object stacks. This is a supplement figure to the synthetic data benchmark shown in Figure 2. Channels 0 and 1 were segmented by Otsu thresholding, and channel 2 was segmented as an occupancy mask. (a–d) Representative benchmark stack shown as channel 0, channel 1, channel 2, and a composite view. (e–h) Corresponding CellColoc masks for the cell, marker, region, and positive-cell outputs. (i–l) Ground truth versus CellColoc estimates for channel-0 counts, channel-1 counts, channel-1-positive channel-0 counts, and channel-2 occupancy. Each line connects the same stack across the two summaries. (m) Mean instance-detection precision, recall, and F1 scores for channels 0 and 1. (n) Mean absolute error summaries across the four benchmark targets. One-to-one matched channel-0 overlap fractions. (p) One-to-one matched channel-0 object areas. (q) One-to-one matched channel-0 roundness values. (r) One-to-one matched channel-0 eccentricity values. Point colors encode stacks.

## Supplementary material

### Details of the microglia experiment

#### Mouse details

The microglia experiment was a small cohort study designed to assess tdTomato reporter expression in *Cx3cr1-creER*^cre/wt^; *Rosa26_tdTom*^fl/wt^ mice with or without tamoxifen induction. The cohort comprised adult C57BL/6J mice pooled across sexes, with a mean endpoint age of approximately 40.6 weeks (range 39.9–41.3 weeks; Table S1). Four mice received tamoxifen and four received vehicle only (Miglyol). The paired regional CA1/CTX comparison shown in the main text used the subset of mice for which both CA1 and CTX image stacks were recorded; two additional mice contributed only single-region acquisitions and were therefore not included in the paired CTX-versus-CA1 testing. All animal experiments were conducted under project license 81-02.04.2023.A102 approved by the competent animal-experimentation authority of the State of North Rhine-Westphalia, Germany.

**Table S1.** Microglia cohort used for the biological experiment. The table lists the animals in the practical-series cohort from which the CA1/CTX imaging subset was drawn. The brain-region column indicates which acquisitions were available for each mouse; only mice with both CA1 and CTX data contributed to the paired regional comparison.

| ID | Sex | Age (weeks) | Treatment | Brain regions imaged | Strain | Genotype |
| --- | --- | --- | --- | --- | --- | --- |
| 22461 | M | 41.3 | Tamoxifen | CA1, CTX | C57BL/6J | *Cx3cr1-creER*<sup>cre/wt</sup>; *Rosa26-tdTom*<sup>fl/wt</sup> |
| 22462 | M | 41.3 | Tamoxifen | CA1, CTX | C57BL/6J | *Cx3cr1-creER*<sup>cre/wt</sup>; *Rosa26-tdTom*<sup>fl/wt</sup> |
| 22463 | M | 41.3 | Vehicle | CA1 | C57BL/6J | *Cx3cr1-creER*<sup>cre/wt</sup>; *Rosa26-tdTom*<sup>fl/wt</sup> |
| 22464 | M | 41.3 | Vehicle | CA1, CTX | C57BL/6J | *Cx3cr1-creER*<sup>cre/wt</sup>; *Rosa26-tdTom*<sup>fl/wt</sup> |
| 22485 | F | 39.9 | Tamoxifen | CA1, CTX | C57BL/6J | *Cx3cr1-creER*<sup>cre/wt</sup>; *Rosa26-tdTom*<sup>fl/wt</sup> |
| 22486 | F | 39.9 | Tamoxifen | CA1, CTX | C57BL/6J | *Cx3cr1-creER*<sup>cre/wt</sup>; *Rosa26-tdTom*<sup>fl/wt</sup> |
| 22487 | F | 39.9 | Vehicle | CA1 | C57BL/6J | *Cx3cr1-creER*<sup>cre/wt</sup>; *Rosa26-tdTom*<sup>fl/wt</sup> |
| 22488 | F | 39.9 | Vehicle | CA1, CTX | C57BL/6J | *Cx3cr1-creER*<sup>cre/wt</sup>; *Rosa26-tdTom*<sup>fl/wt</sup> |

#### Mouse preparation

To induce Cre recombinase activity in microglia, tamoxifen (Sigma-Aldrich, St. Louis, Missouri, USA) was administered intraperitoneally at 0.1 mg/g body weight on five consecutive days. Tamoxifen was dissolved in Miglyol 812 neutral oil (Caesar & Loretz GmbH, Hilden, Germany) by sonication before injection. Vehicle-control mice received Miglyol alone on the same schedule. Animals were perfused ten days after the final injection by transcardial perfusion with phosphate-buffered saline (PBS) followed by paraformaldehyde (PFA) fixation overnight. Brains were collected and sectioned coronally at 100 *µ*m thickness using a vibratome. Two coronal slices per mouse were selected at approximately Bregma −1.5 to sample the hippocampal CA1 region and the somatosensory cortex. Free-floating sections were immunostained over two days with rabbit anti-Iba1 (Wako, 019-19741; 1:1000), Alexa Fluor 488 goat anti-rabbit secondary antibody (Invitrogen, A11008; 1:500), and DAPI (Sigma-Aldrich, D9542; 1:10000 during the final 5 min of staining), and were then mounted in Dako mounting medium. In this experiment, tdTomato marked the recombined CX3CR1-positive microglial population, Iba1 served as an independent microglia/macrophage-lineage marker, and DAPI was used primarily for anatomical orientation and quality control.

#### Imaging

All analyzed image stacks were acquired as three-channel Zeiss CZI files on a Zeiss LSM 800 inverted laser-scanning confocal microscope equipped with a Plan-Apochromat 20x/0.8 M27 air objective. The recorded fluorescence channels corresponded to tdTomato/mCherry, Alexa Fluor 488 for Iba1, and Alexa Fluor 405 for DAPI. Stacks were acquired at 1024 *×* 1024 pixels over 30 optical sections with 1 *µ*m z spacing and physical sampling of 0.6239 *µ*m in *x* and *y*, corresponding to an in-plane field of view of approximately 638.9 *×* 638.9 *µ*m per stack.

#### Statistics

The regional microglia comparison in Figure 3 used mouse as the experimental unit. When more than one stack was available for the same mouse and brain region, region-specific summary values were averaged first so that each matched mouse contributed one CTX value and one CA1 value per endpoint. Mice represented in only one of the two regions were excluded from the paired comparison, which yielded six matched CA1/CTX pairs in the analyzed image collection.

Normality was assessed on the paired CA1–CTX differences for each endpoint using the Shapiro–Wilk test [26]. If normality was not rejected at *α* = 0.05 (that is, if the Shapiro–Wilk *p* value exceeded 0.05), a two-sided paired *t*-test was used. Otherwise, a two-sided Wilcoxon signed-rank test was applied. In the edge case of exactly zero paired differences across all matched mice, the analysis script returned *p* = 1.0 by construction. All *p* values reported for the regional comparisons in Figure 3 were obtained from this mouse-level paired analysis. No multiple-testing correction was applied to the Figure 3 comparisons shown in the present preprint.

For the treatment-sensitivity analysis shown in Figure 3 g)–j), mouse was again used as the experimental unit. Repeated scans were first averaged within each mouse and region. The available regional values were then averaged within mouse to obtain one global treatment-level value per endpoint and animal. This allowed mice with only one available region to contribute to the treatment-sensitivity analysis while keeping the statistical unit at the mouse level, yielding four vehicle-treated and four tamoxifen-treated mice. Vehicle-only and tamoxifen-treated animals were compared with two-sided Mann–Whitney U tests. These comparisons were used as a small-cohort sensitivity check for treatment-associated shifts in the key tdTomato/Iba1 readouts and average tdTomato cell brightness rather than as the primary biological endpoint.

All statistical calculations were carried out in Python 3.12.13 using NumPy 2.4.6 [27] and SciPy 1.18.0 [28]. The normality and paired-comparison tests were evaluated with functions from the `scipy.stats` module.

